## Supplementary Figures for "CD131 Contributes to Ulcerative Colitis Pathogenesis by Promoting Macrophage Infiltration"

by Zhiyuan Wu *et al.*

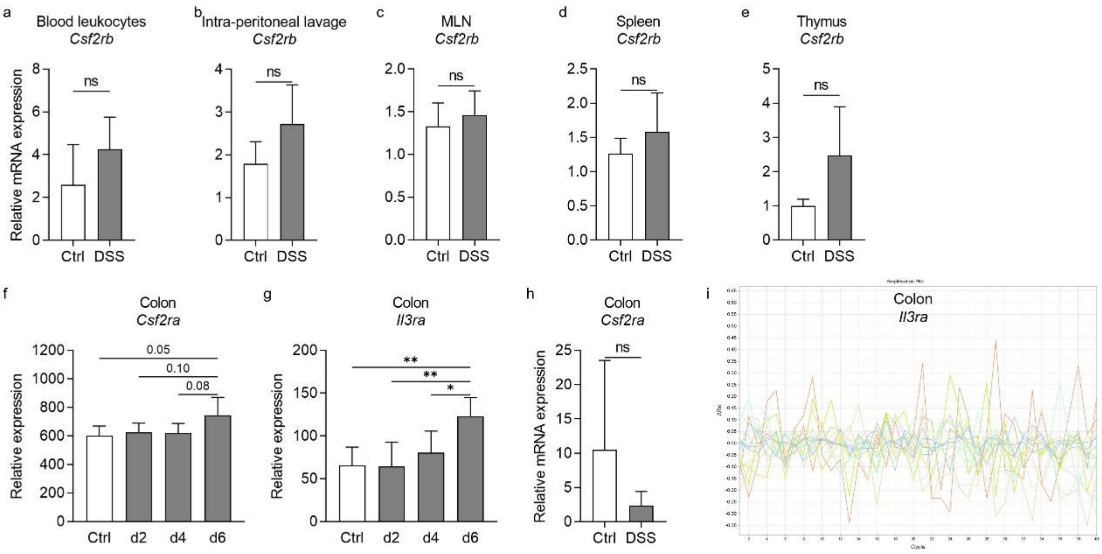


Fig S1. (a-e) Relative *Csf2rb* mRNA expression levels in blood leukocytes (a), intra-peritoneal lavage cells (b), mesenteric lymph nodes (MLN, c), spleen (d) and thymus (e) of control and DSS-treated wt mice. (f-g) Gene expression dataset shows the change of relative *Csf2ra* and *Il3ra* gene expression levels in murine colon tissues with time after DSS administration. (h) Relative *Csf2ra* mRNA expression level in colon tissues of control and DSS-treated wt mice. (i) Amplification plot from Applied Biosystems 7500 Fast Real-Time PCR System showing that *Il3ra* mRNA expression in colon tissues was not detectable.

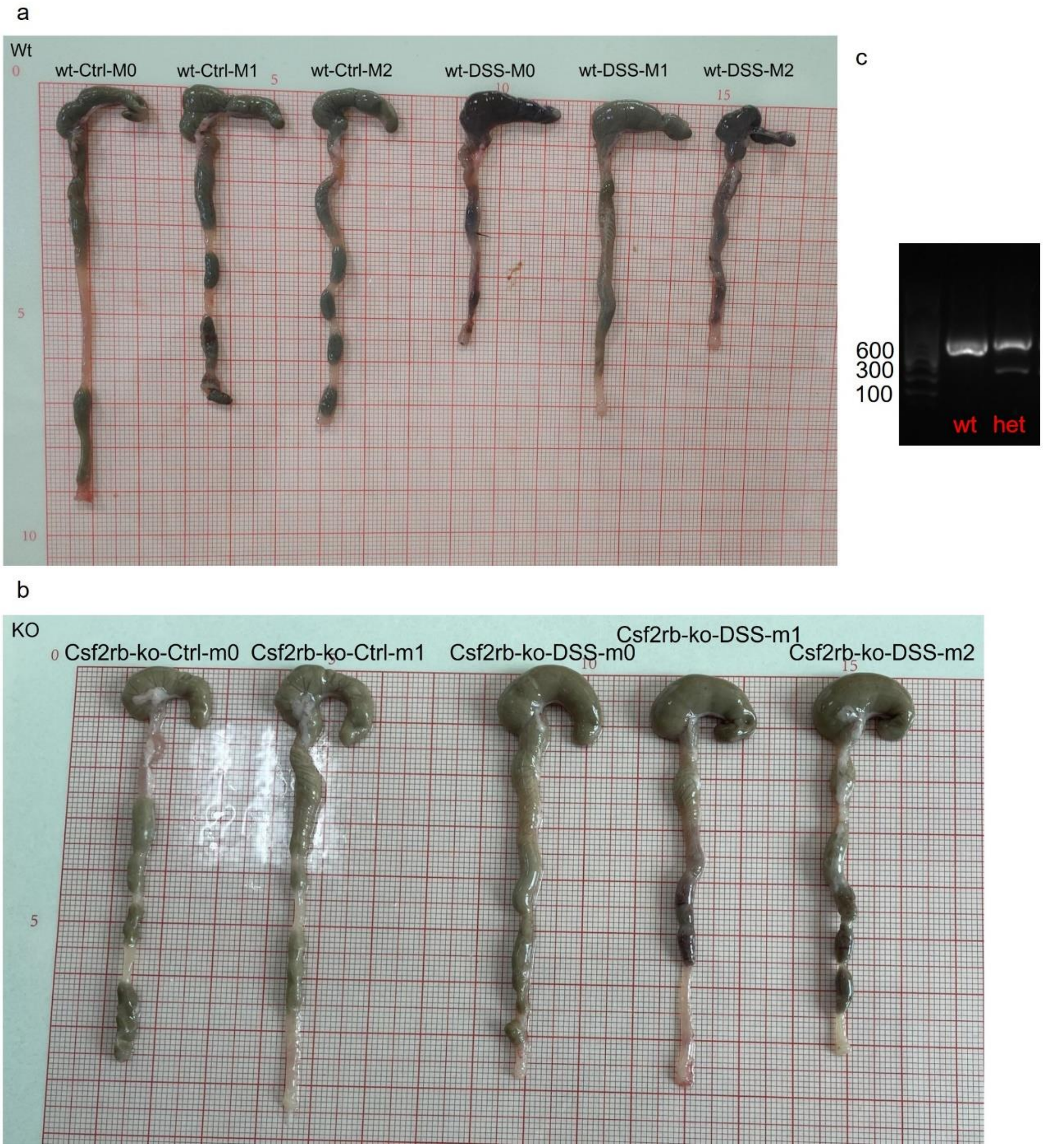


Fig S2. (a-b) Exemplary photos of colons sampled from wt and CD131-deficient mice that were treated with normal drinking water (Ctrl) or DSS. (c) An exemplary graph of 1.5% agarose gel electrophoresis showing wt and heterozygous knock-out (het) genotypes on murine tail biopsies genotyping.

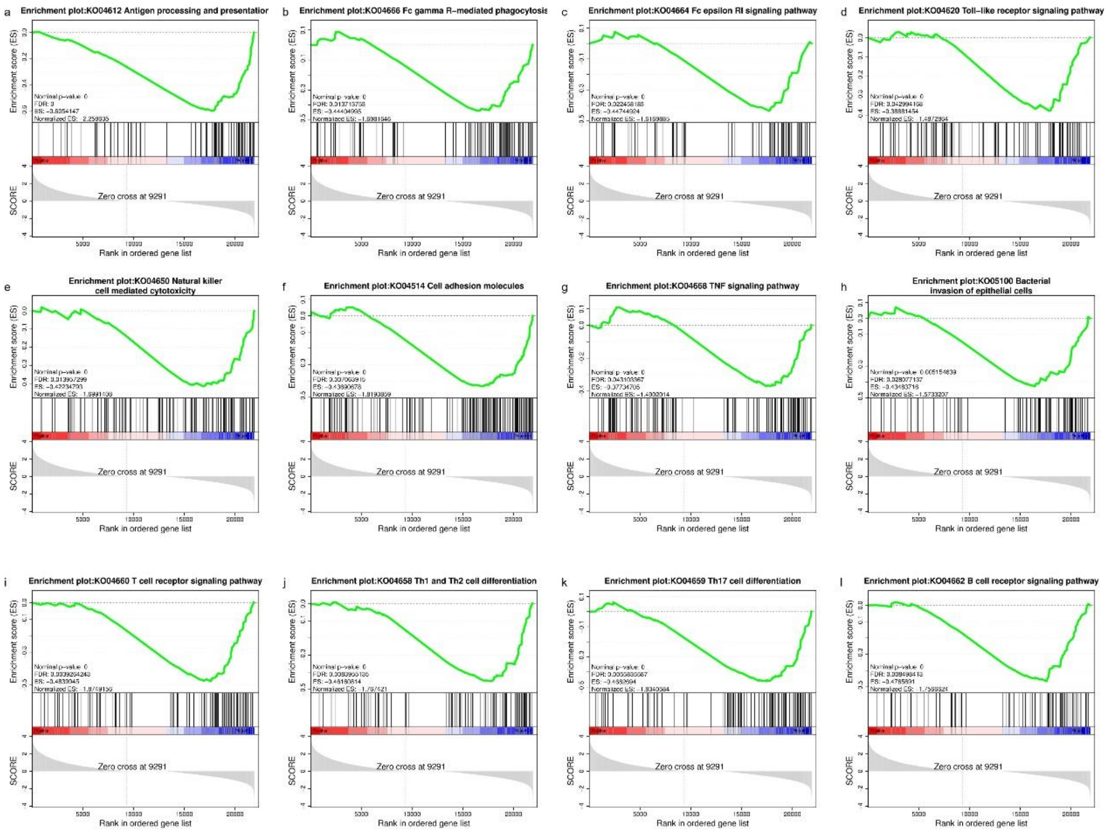


Fig S3. GSEA of colon tissues between wt and CD131-deficient mice showing that several innate and acquired immune-related pathways, including (a) antigen processing and presentation, (b) FcγR-mediated phagocytosis, (c) FcεRI signaling pathway, (d) Toll-like receptor signaling pathway, (e) natural killer cell-mediated cytotoxicity, (f) cell adhesion molecules, (g) TNF signaling pathway, (h) bacterial invasion of epithelial cells, (i) T cell receptor signaling pathway, (j) Th1 and Th2 cell differentiation, (k) Th17 cell differentiation and (l) B cell receptor signaling pathway, were enriched in wt mice.

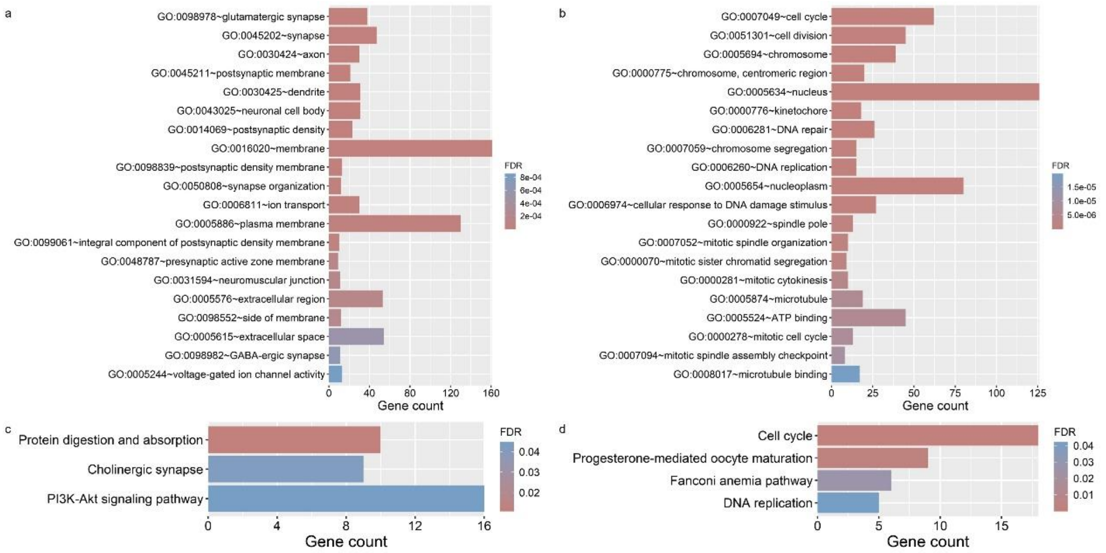


Fig S4. (a-b) Gene Ontology (GO) enrichment analysis of the up- (a) and down-regulated (b), intersected, DEGs. (c-d) Kyoto Encyclopedia of Genes and Genomes (KEGG) enrichment analysis of the up- (c) and down-regulated (d), intersected, DEGs.

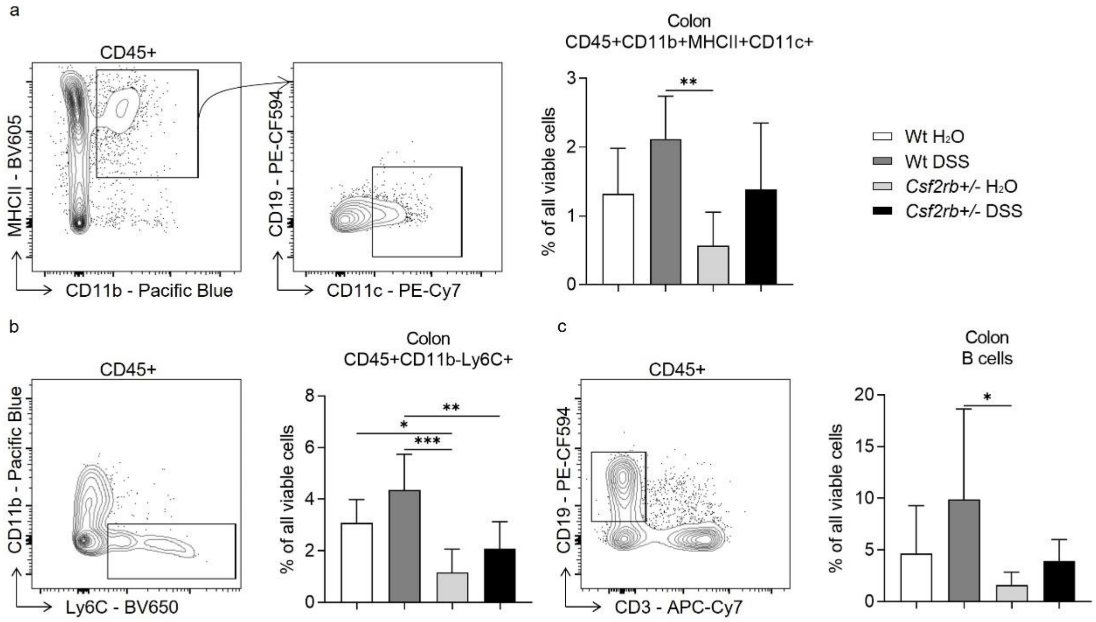


Fig S5. (a) Exemplary graphs showing the gating strategy for identifying CD45⁺CD11b⁺MHCII⁺CD11c⁺ cells on multi-color flow cytometry and their relative cell number, normalized to all viable cells, in the colonic tissues of wt and CD131-deficient mice treated with normal drinking water or DSS. (b) An exemplary graph showing the gating strategy for identifying CD45⁺CD11b⁻Ly6C⁺ cells on multi-color flow cytometry and their relative cell number, normalized to all viable cells, in the colonic tissues of wt and CD131-deficient mice treated with normal drinking water or DSS. (c) An exemplary graph showing the gating strategy for identifying CD45⁺CD19⁺CD3⁻ B cells on multi-color flow cytometry and their relative cell number, normalized to all viable cells, in the colonic tissues of wt and CD131-deficient mice treated with normal drinking water or DSS.

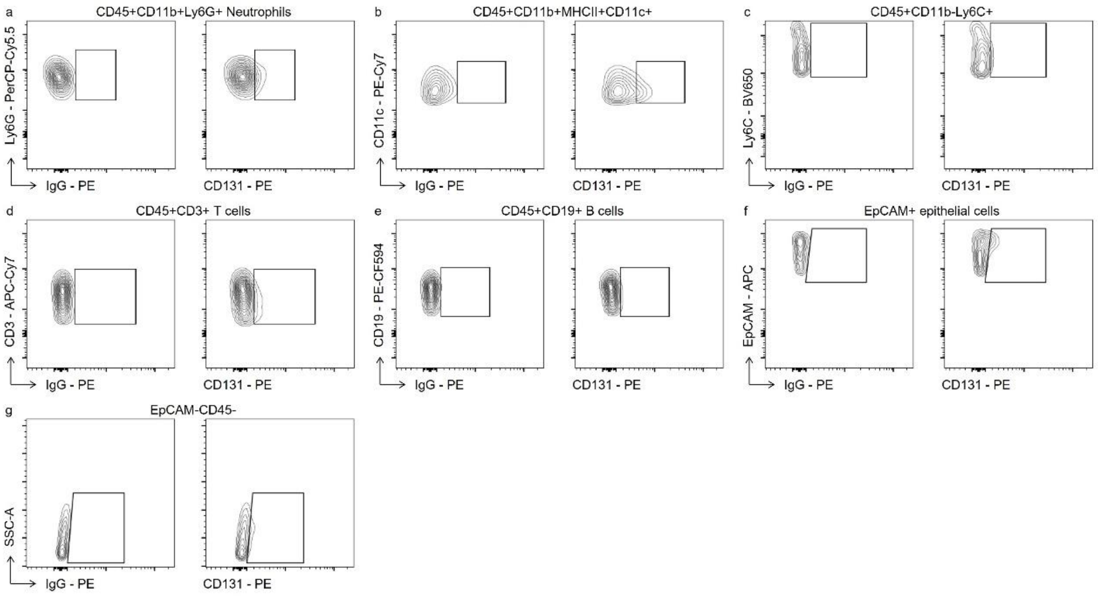


Fig S6. Exemplary graphs showing the gating strategy for identifying (a) $CD131^{+}CD11b^{+}Ly6G^{+}$ neutrophils, (b) $CD131^{+}CD45^{+}CD11b^{+}MHCII^{+}CD11c^{+}$ cells, (c) $CD131^{+}CD45^{+}CD11b^{-}Ly6C^{+}$ cells, (d) $CD131^{+}CD45^{+}CD3^{+}$ T cells, (e) $CD131^{+}CD45^{+}CD19^{+}$ B cells, (f) $CD131^{+}EpCAM^{+}$ epithelial cells and (g) $CD131^{+}EpCAM^{-}CD45^{-}$ cells.

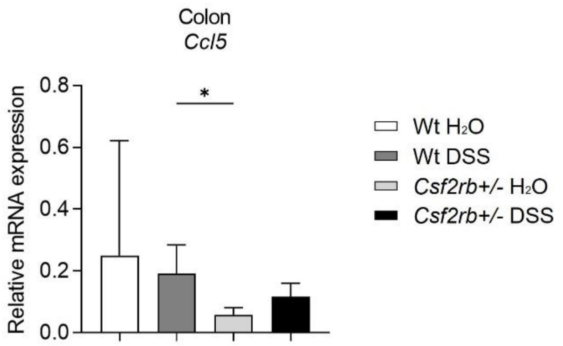


Fig S7. Relative *Ccl5* mRNA expression levels in colon tissues of wt and CD131-deficient mice treated with normal drinking water or DSS.
