## Supplementary material for "CD131 Contributes to Ulcerative Colitis Pathogenesis by Promoting Macrophage Infiltration": Table S7

**Table S7. Demographic and clinical data of ulcerative colitis patients.**

| **#** | **Age (yr)** | **Gender** | **Hx (mo)** | **Smoking** | **Temp (°C)** | **Pulse (per min)** | **Stool freq (per day)** | **Blood in stool** | **Extent (Montreal)** | **WBC (x10^9/l)** | **Hb (g/dl)** | **ESR (mm/h)** | **Mucosal appearance at endoscopy (Mayo)** |
| --- | --- | --- | --- | --- | --- | --- | --- | --- | --- | --- | --- | --- | --- |
| 1 | 44 | F | 216 | Never | 36.5 | 88 | 2 | Fobt+ | E3 | 5.22 | 9.2 | na | Severe |
| 2 | 62 | M | 240 | Current | 36.5 | 80 | 4-5, to >10 | Fobt+ | E2 | 11.59 | 12.4 | na | Severe |
| 3 | 50 | M | 240 | Never | 36.3 | 114 | >20 | Visible | E3 | 5.2 | 7.7 | na | Severe |
| 4 | 53 | M | 240 | Never | 36.8 | 78 | 2-3 | Visible | E3 | 7.28 | 12.8 | na | Moderate |
| 5 | 44 | M | 84 | Never | 36.0 | 80 | 2-3 d/time | Fobt+ | E3 | 5.6 | 15.7 | na | na |
| 6 | 53 | F | 240 | Never | 36.9 | 82 | 7-8 | Visible | E2 | 12.7 | 9.4 | 82 | Severe |
| 7 | 51 | F | 240 | Never | 36.5 | 70 | >10 | Visible | E3 | 10.1 | 9.3 | na | Severe |
| 8 | 58 | M | 36 | Never | 36.5 | 74 | na | Visible | E3 | 7.67 | 9.3 | na | Severe |
| 9 | 35 | F | 48 | Never | 36.8 | 90 | 8-10 | Visible | E3 | 6.05 | 7.6 | na | Severe |
| 10 | 41 | M | 168 | Never | 36.5 | 100 | 15 | Visible | E3 | 28.54 | 7.2 | 48 | Severe |
| 11 | 52 | F | 216 | Never | 36.5 | 76 | >10 | Visible | E3 | 8.21 | 9.2 | 12 | Severe |
| 12 | 54 | F | 312 | Never | 36.5 | 70 | >20 | Fobt+ | E2 | 5.05 | 7.5 | 2 | Severe |
| 13 | 55 | M | 12 | Former | 36.4 | 74 | 3 | Visible | E3 | 12.52 | 7.8 | na | Severe |
| 14 | 80 | F | 12 | Never | 36.5 | 80 | 4-5 | Visible | E3 | 7.17 | 10.5 | na | Severe |
| 15 | 57 | F | 60 | Never | 36.2 | 70 | na | Visible | E3 | 6.3 | 7.7 | na | na |
| 16 | 70 | M | 12 | Never | 36.5 | 80 | >10 | Visible | E3 | 6.5 | 9.1 | 40 | Severe |
| 17 | 54 | M | 5 | Former | 36.0 | 82 | 5-6 | Visible | E3 | 12.9 | 8 | 75 | Severe |
| 18 | 50 | M | 12 | Former | 36.5 | 102 | 10 | Visible | E3 | 7.81 | 9.1 | 65 | Severe |
| 19 | 58 | F | 36 | Former | 36.5 | 75 | 1-2 | Visible | E3 | 3.56 | 8 | 64 | Severe |
| 20 | 62 | M | 120 | Former | 36.5 | 70 | 6-7 | Visible | E3 | 11.24 | 10.2 | 61 | Severe |
| 21 | 36 | M | 36 | Never | 37.2 | 90 | 3-4, to 6 | Visible | E3 | 13.54 | 6.7 | 76 | Severe |
| 22 | 42 | F | 96 | Never | 36.5 | 80 | 5-6 | Fobt+ | E3 | 4.3 | 8.8 | 2 | Severe |
| 23 | 52 | M | 5 | Former | 36.5 | 78 | 7-8 | Visible | E3 | 4.71 | 6.2 | na | na |
| 24 | 67 | F | 36 | Never | 36.5 | 84 | 7-8 | Visible | E3 | 5.68 | 9.4 | 14 | Severe |
| 25 | 51 | F | 12 | Never | 36.5 | 80 | >20 | Visible | E3 | 3.8 | 9.1 | 18 | Severe |
| 26 | 56 | M | 120 | Never | 36.5 | 96 | >10 | Visible | E3 | 5.2 | 8.3 | 10 | Moderate |
| 27 | 50 | M | 1 | Current | 36.5 | 92 | 20-30 to 5-6 | Visible | E3 | 3.6 | 8.2 | 13 | Moderate |
| 28 | 65 | M | 2 | Former | 36.5 | 76 | 6-7 to 20 | Visible | E3 | 3.7 | 8.3 | 7 | Severe |
| 29 | 48 | F | 1 | Never | 36.7 | 66 | na | Visible | E3 | 4.9 | 13.1 | na | Moderate |

ESR: erythrocyte sedimentation rate; Fobt: fecal occult blood test; na: not available (data missing from clinical registry).
